# Tick-borne pathogens identified in *Hyalomma marginatum* (Acari: Ixodidae) collected from various vertebrate hosts in the South of France

**DOI:** 10.1101/2024.03.01.582933

**Authors:** Bernard Célia, Pollet Thomas, Galon Clémence, Joly Kukla Charlotte, Cicculli Vincent, Falchi Alessandra, Grech-Angelini Sebastien, Poli Paul-Eric, Bastien Matthieu, Combes Benoit, Moutailler Sara, Holzmuller Philippe, Vial Laurence

## Abstract

*Hyalomma marginatum* is a common ectoparasitic tick of ungulates, lagomorphs, insectivores, ground-foraging birds, observed in Corsica for decades, but whose permanent establishment in mainland France is very recent. This species is known to be one of the main vectors of the Crimean-Congo Hemorrhagic Fever virus, but also of various parasitic, bacterial or viral pathogens. In this study, we investigated the molecular infection rates of numerous tick-borne pathogens in *H. marginatum* ticks mainly sampled on horses, and occasionally on other animal species, from the French Mediterranean rim and Corsica between 2016 and 2020. In total, 1, 195 DNA and RNA purified from individual ticks or pools of ticks were screened for 26 microbial genera or species (viruses, bacteria and parasites), using a high-throughput microfluidic real-time PCR system (BioMark™ dynamic array system, Standard Biotools). For individual ticks and pooled ones, respectively, the most prevalent tick-borne microorganisms were *Francisella*-like endosymbionts at 97.0% and 96.8%, followed by *Rickettsia aeschlimannii* (76.4% and 96.4%), *Theileria* spp. and *Theileria equi* (3,5% and 0%; 1,9% and 5,8%), *Anaplasma phagocytophilum* (3.7% and 6.7%), and West Nile virus (0.1% and 0.4%). *Babesia occultans* (0.9%), *Ehrlichia minasensis* (0.3%), and *Coxiella*-like endosymbionts (0.1%) were only detected in individual ticks. Our study provides an overview of the diversity of microorganisms and tick-borne pathogens detected in the invasive tick *H. marginatum* in Corsica and the continental departments of the Mediterranean rim. Our study opens up new research perspectives on the epidemiology of tick-borne pathogens carried by *H. marginatum* and on the associated public and veterinary health risks.

## Introduction

Worldwide, ticks are considered to be the first vectors of pathogens responsible for animal diseases and second behind mosquitoes for those affecting humans (Jongejan et Uilenberg 2004; Colwell, Dantas-Torres, et Otranto 2011). In France, about forty species of ticks are identified on the territory, most of them transmitting pathogens responsible for important public and veterinary health problems (Boulanger et McCoy 2017; PÉREZ-EID 2007). This specifically includes *Ixodes ricinus*, the primary vector for the agents of Lyme borreliosis, *Borrelia burgdorferi* sensu lato, as well as ticks from the genera *Dermacentor* or *Rhipicephalus* responsible for the transmission of equine piroplasmosis (Gray 1991; Nadal, Bonnet, et Marsot 2022). Recently, Vial et al. (2016) provided strong evidence for the presence of reproducing populations of the exotic tick *Hyalomma marginatum* in parts of southern France. While this tick species has been present in Corsica (a Mediterranean island of southern France) for several decades, it seems to be now established on the Mediterranean coastline of mainland France in many areas such as Pyrénées-Orientales, Aude, Hérault, Gard, Ardèche, Drôme and Var (Bah et al. 2022). This tick specieshas been also detected since many years in Spain (Palomar et al. 2016) and Italy (Satta et al. 2011), showing a possible continuum for the sanitary risks associated to *H. marginatum* in southwestern Europe. As a “generalist” tick, *H. marginatum* presents a large panel of vertebrate hosts, ranging from lagomorphs and birds for the immature stages to large domestic and wild ungulates for the adult ones (Bernard et al., 2022). It is a two-host tick, meaning that larvae and nymphs feed on the same individual, while the nymph detaches and moults into an adult that seeks out a new host for blood feeding and mating. This diversity of hosts expands the pathways for tick introduction via natural or human-driven animal movements and is believed to increase opportunities for pathogen transmission and possible spillover. Ticks of the genus *Hyalomma* are involved in the transmission of many human and animal pathogens in Eurasia and Africa (Bonnet et al. 2023). They can in particular carry parasites and bacteria such as *Theileria* spp., *Babesia* spp. and *Anaplasma* spp., responsible for equine piroplasmosis or bovine anaplasmosis, with often non-specific clinical signs (e.g. hyperthermia, asthenia, abortion…) in cattle or sheep (Wise et al. 2013, Rothschild et al. 2013, Ristic et al., 1981). Other infectious bacteria such as *Coxiella burnetii*, and *Rickettsia* spp. including *R aeschlimannii* (Bakheit et al. 2012; Kilicoglu et al. 2020; Orkun 2022), or viruses such as Bhanja (Genus Bandavirus) and West Nile (Genus Flavivirus) (Formosinho et Santos-Silva 2006; Kolodziejek et al. 2014), can be carried by *H. marginatum*. Additionally, Orthonairoviruses such as the emerging and highly contagious Crimean Congo Hemorrhagic Fever virus (CCHFV) (Bakheit et al. 2012) can be transmitted by this tick spcies. In the Mediterranean basin, two species of *Hyalomma* (*H. marginatum* and *H. anatolicum*) are used to be considered as the main vectors of CCHFV (Perveen et Khan 2022). However, viral RNA has been recently detected in *H. lusitanicum* collected from deer and wild boar in Spain, in higher proportion than in *H. marginatum*, suggesting major role of this additional tick in CCHFV transmission in this country (Moraga-Fernández et al. 2021). In France, based on our knowledge on local tick communities, we considered *H. marginatum* as the main candidate tick species likely to transmit CCHFV (Bernard et al. 2022). In 2023, this was confirmed by the first detection of CCHFV in H. marginatum ticks from Pyrénées-Orientales (Bernard, 2024). Considering the large geographic distribution of *H. marginatum*, from Spain to North Africa and India (Fernández-Ruiz et Estrada-Peña 2021), this questions about local adaptations in the abilities developed by this tick species to maintain and transmit all, or part of the pre-cited pathogens.

The objective of this study was thus to describe the panel of pathogens and/or endosymbiotic microorganisms in *H. marginatum* ticks locally collected in the South of France, mainland and Corsica island. This work was part of a national surveillance program developed a few years ago to assess the risk of emergence of exotic diseases including tick-borne diseases in France, under the aegis of the French Ministry of Agriculture. In Corsica, first investigations were recently conducted on tick species collected from different hosts, especially cattle, and reported in *H. marginatum* high prevalence of *Francisella*-like endosymbionts, *R. aeschlimannii* and in a less extend *Candidatus Ri. Barberiae*, as well as the presence of *Anaplasma marginale* and *A. phagocytophylum* (V. Cicculli et al. 2019; Grech-Angelini et al. 2020). These studies also allowed the detection of new viruses in *H. marginatum*, for example a Pseudocowpox virus of which the zoonotic potential remains unknown (V. Cicculli et al. 2019). On mainland Francewhere *H. marginatum* was not yet established only 15 years ago, such investigation has never been done. Keeping in mind that genomic detection of pathogens does not imply a mandatory vector competence of the tick for the considered microorganisms (Bonnet et al. 2023), this study aimed to assess potential vector competence by combining pathogen detection with an understanding of *H. marginatum*’s host preferences and the susceptibility of these hosts to the identified pathogens. The aim was to know whether *H. marginatum* ticks arrived with their native associated pathogens usually described in endemic Mediterranean areas including Corsica, and may result in the introduction of new pathogens in France. Another alternative would be that *H. marginatum* can become a new vector for local French tick-borne pathogens already present on the territory, with the risk of changing the local transmission dynamics of such pathogens due to its distinct seasonal activity from other local tick species.

## Methods

### Study area, period and hosts for tick collection

Among all the *H. marginatum* ticks analyzed in this study, 1,711 came from a large-scale tick sampling that was conducted on horses (considered as the likely host of *H. marginatum* adult stages), during spring (the activity period of *H. marginatum* adult stages), in southern France (mainland and Corsica), between 2016 and 2020. This was carried out to determine and explain the distribution of the newly established tick *H. marginatum* on the French territory (Bah et al. 2022). Eight French departments forming a link from Spain to Italy, which are two countries where *H. marginatum* is known to be already present, plus the two departments of Corsica were sampled. *H. marginatum* ticks were detected in 56 and 38 equestrian structures in mainland France and in Corsica, respectively. On the mainland, a reference site in Pompignan, where *H. marginatum* is abundant and monitored for its dynamics of infestation on horses, was also sampled each year and 173 ticks of the 1,711 ticks came from this site. In Corsica, each year, the whole island was visited whereas on the mainland, most of *H. marginatum* were obtained in 2017, except for northern Gard and Ardèche that were investigated in 2019 (Figure 1). Among the 1,711 ticks, 900 collected in Corsica in 2019 and 2020 were grouped into 222 pools of 1 to 10 individuals according to different parameters of sex or hosts (Cicculli et al., 2022) and 811 were treated individually.

**Figure 1:**
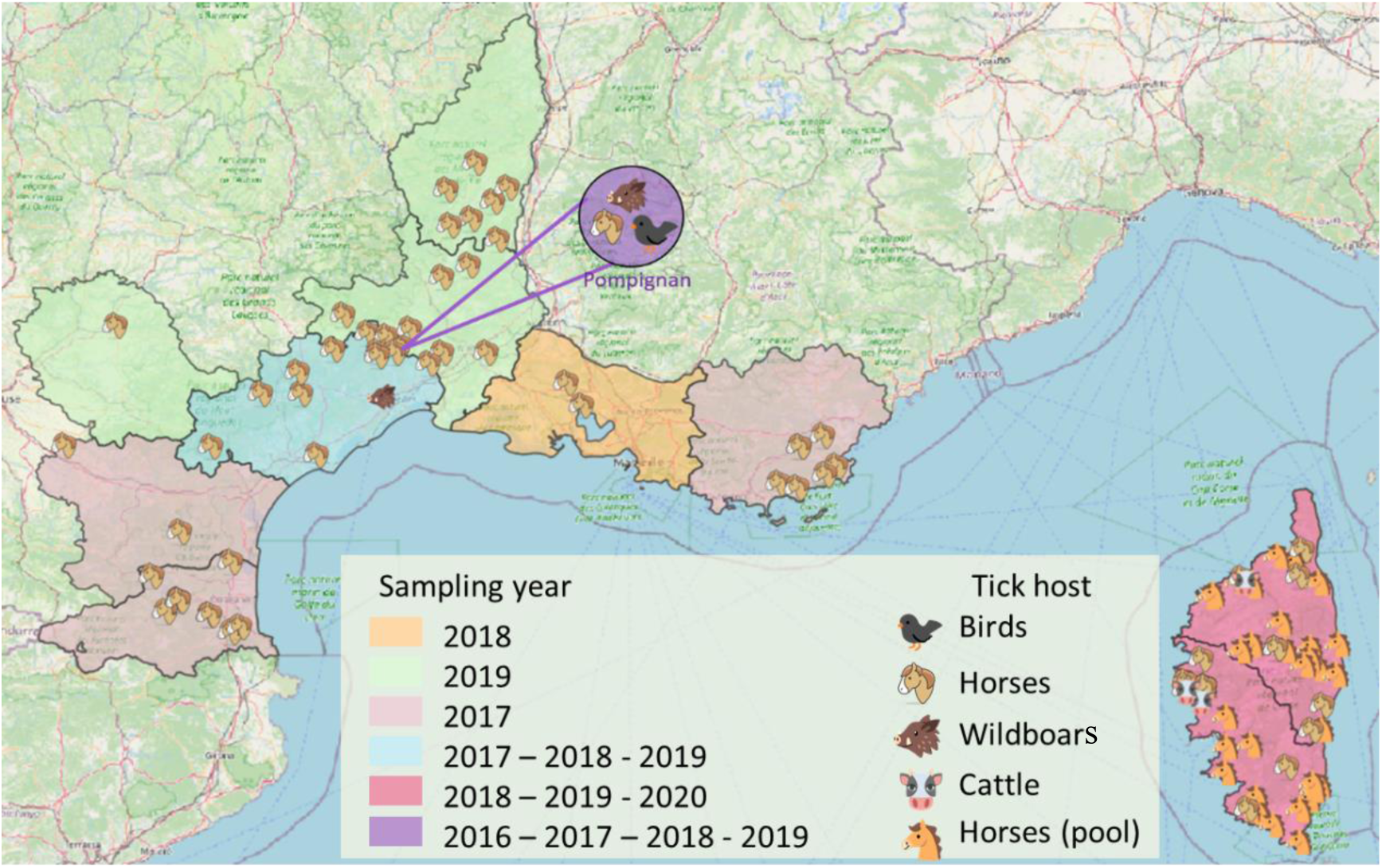
Study area overview : geographic distribution and host diversity. On this map, sites are represented by the host species from which the ticks were collected (horses, cattle, wildboars and birds).

**Figure 2:**
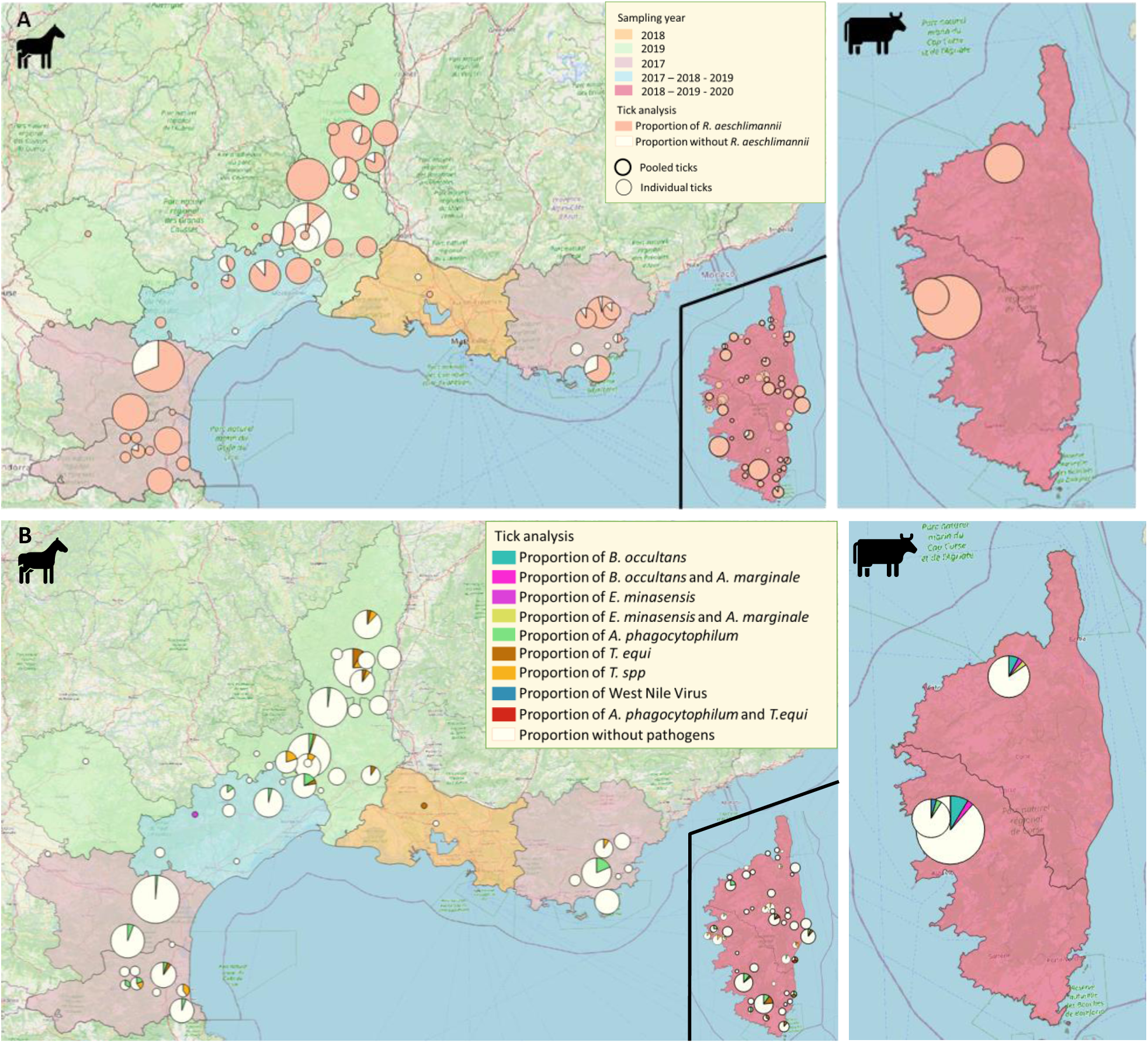
Proportions of *Rickettsia aeschlimannii* infections (A) and other microorganisms (B) according to geographical sampling sites in *H. marginatum* ticks collected on horses (on the left) and cattle (only in Corsica) (on the right). The size of the circles is proportional to the number of ticks collected per sampling site, the color of the department (in A and B) corresponds to the samplig year.

In Corsica, 118 additional ticks were collected from cattle examined in three farms located in the northern part of the island, in 2019.

Additionally, 7 and 37 *H. marginatum* ticks were obtained by punctual samplings in Hérault and Gard on wild boars in 2016 and birds in 2017, respectively. Ticks from birds were all immature stages, and were mostly associated to the resident avian fauna of Pompignan (Vial et al. 2016). Thus, taking all host species together, we worked on 973 ticks treated individually and 222 tick pools (1,195 samples).

During any tick samplings, animals were examined on the whole body and all tick species were collected. Back from the field, they were characterized under binocular magnifiers using morphological identification keys (PÉREZ-EID 2007), and sorted according to species, locality, host, gender (male/female), developmental stage (larva/nymph/adult), and engorgement status. They were then frozen at −80°C for later molecular analysis. As this study focused on sanitary risks associated to the newly established *H. marginatum*, only specimens from this species were screened for microorganisms.

### DNA and RNA extraction

As we looked for DNA and RNA of microorganisms in all tick samples, their treatment for molecular analysis was the same.

After a first washing in bleach for 30 sec followed by three times in distilled water for 1 min, ticks were crushed individually with three steel beads (2.8mm, OZYME, France) in 300µL DMEM (Eurobio Scientific, France) supplemented with 10% fetal calf serum (Dutscher, France), using the homogenizer Precellys® 24 Dual (Bertin, France) at 5500 rpm for 40s. The processing was different for samples of horses coming from Corsica as, prior to be crushed, ticks were pooled by up to 10 individuals per pool, according to locality and developmental stage (Vincent Cicculli et al. 2022).

DNA was extracted using 100μl of the homogenate of crushed ticks according to the Genomic DNA from tissue kit (Macherey-Nagel, Germany) following the manufacturer’s instructions. Total DNA per sample was eluted in 50 μl of rehydration solution and stored at −20◦C until further use (Michelet et al, 2014).

Total RNA was extracted from 100µl of the homogenate of crushed ticks using the NucleoSpin RNA kit (Macherey-Nagel, Germany) according to the manufacturer’s instructions. Total RNA per sample was eluted in 50 μl of RNA free water and stored at −80◦C until further use (Gondard et al., 2018).

### Detection of tick-borne microorganisms’ DNA or RNA

The Reverse Transcription Master Mix kit (Standard Biotools, USA) was used on all samples to obtain cDNAs from RNA viruses. The reaction was performed in 5μL of final volume, including 1μL of Reverse Transcription Master Mix, 3μL of RNase-free ultrapure water, and 1μL of extracted RNA. A thermal cycler (Eppendorf, Germany) performed the RT cycles following the program: 25°C for 5 min, 42°C for 30 min and a final step at 85°C for 5 min for a total duration of 40 min.

A pre-amplification step was performed to pre-amplify the DNA/cDNA to increase the signal of the genetic material of the microorganisms compared to those of the tick. The PreAmp Master Mix kit (Standard Biotools, USA) was used. A 0.2X pool containing the primer pairs of the targeted pathogens was made upstream, to obtain a final primer concentration of 200 nM. For one sample, the mix was composed of 1 μL of PreAmp Master mix, 1.25 μl of the 0.2X pool, and 1.5 μl of ultra-pure water. Then 1.25μL of sample (DNA/cDNA mix) was added. A negative control (ultra-pure water) was added to each plate. PCR was then performed using a thermal cycler with the following program: 95°C for 2min, 14 cycles of two steps of 95°C for 15s and a final step of 4min at 60°C.

Then, for the real time microfluidic PCR, twenty-four sets of primers and probes were used to detect predefined microorganisms specifically targeted for this study (Table 1) (Michelet et al. 2014; Gondard et al. 2018). The BioMark™ real-time PCR system (Standard Biotools, USA) was used for high throughput microfluidic real-time PCR amplification using 48.48 dynamic arrays (Standard Biotools). These chips dispense 48 PCR mixes and 48 samples into individual wells, after which on-chip microfluidics assemble PCR reactions in individual chambers prior to thermal cycling resulting in 2,304 individual reactions (Michelet et al. 2014). In a single experiment, 47 individual ticks or tick pools samples (and one negative control) can be tested.

**Table 1:** Tick-borne pathogens and endosymbionts targeted by the chip of the high-throughput microfluidic real-time PCR system developed specifically for the study or for specific RT-qPCR (CCHFV).

| Genus / Family | Species |
| --- | --- |
| *Bacteria* |  |
| *Anaplasma* | *A. marginale*, *A. phagocytophilum*, *Anaplasma* spp. |
| *Bartonella* | *B. henselae*, *Bartonella* spp. |
| *Borrelia* | *B. miyamotoi*, *Borrelia* spp. |
| *Coxiella* | *Coxiella burnetii* and *Coxiella*-like endosymbionts |
| *Ehrlichia* | *Ehrlichia* spp. |
| *Francisella* | *F. tularensis*, *Francisella*-like endosymbionts |
| *Rickettsia* | *R. aeschlimannii*, *Rickettsia* spp. |
| *Protozoan* |  |
| *Babesia* | *B. caballi* |
| *Theileria* | *T. equi*, *T. annulata*, *Theileria* spp. |
| *Viruses* |  |
| *Bunyaviridae* | Nairobi sheep disease virus |
| *Flaviviridae* | West Nile Virus, Alkhurma virus |
| *Nairoviridae* | Hazara virus, CCHFV |
| *Orthomyxoviridae* | Thogoto virus, Dhori virus |
| *Phenuiviridae* | Bhanja virus |

In order to develop the a priori approach to detect pathogens in ticks using real time microfluidic PCR, we first generated a list of microorganisms (TBPs and endosymbionts) (Table 1) with the highest probability of being transmitted or carried by *H. marginatum*, according to data mining of the available literature. From two studies *(Bakheit et al. 2012; Bonnet et al. 2022)*, we identified several potentially *H. marginatum*-borne pathogens: protozoa such as *B. caballi* or *T. annulata*, bacteria such as *R. aechlimannii* and finally viruses such as Thogoto, Dhori and Bhanja viruses (RNA for these viruses has already been detected in ticks), in addition to CCHFV. We then identified microorganisms from scientific literature that mentioned the detection of pathogens in *H. marginatum*, which could be transmitted by this tick or just reflect the infectious status of the animal host on which the tick was collected, as shown in Corsica or other regions of the world: *A. phagocytophilum*, *A. marginale (Grech-Angelini et al. 2020)*, *E. minasensis (Vincent Cicculli et al. 2019)*, *T. equi (Rocafort-Ferrer et al. 2022)*, the West Nile Virus *(Formosinho et Santos-Silva 2006; Kolodziejek et al. 2014)*, the symbiotic form of *Coxiella burnetii* and *Francisella tularensis* : *Coxiella*-like endosymbiont CLE and *Francisella*-like endosymbiont FLE *(Kilicoglu et al. 2020; Selmi et al. 2019; Buysse et al. 2021; Demir et al. 2020)*. Then, some pathogens known to be transmitted by ticks of the genus *Hyalomma* but not necessarily *H. marginatum*, or even by other tick genera but with a likelihood to replicate in *Hyalomma* ticks, were also added to the chip: Alkhurma virus transmitted by *H. rufipes (Horton et al. 2016; Hoffman et al. 2018)*, as well as Nairobi sheep disease and Hazara viruses that are genetically close to CCHFV (Garrison *et al., 2020)*, *Borrelia miyamotoi* and *Bartonella henselae* that circulate in France in other tick species or even other arthropoda such as fleas *(Cosson et al. 2014; Grech-Angelini et al. 2020)*.

To confirm our results of positive detection of pathogens by the microfluidic chip in field samples, specific PCRs were performed individually on some samples with the following primers (Table 2). Conventional PCRs using primers targeting genes or regions other than those of the BioMark™ system were used. Amplicons were sequenced by Eurofins MWG Operon (Germany), and then assembled using BioEdit software (Ibis Biosciences, Carlsbad). An online BLAST (National Center for Biotechnology Information) was used to compare results with published sequences listed in GenBank sequence databases.

**Table 2:**
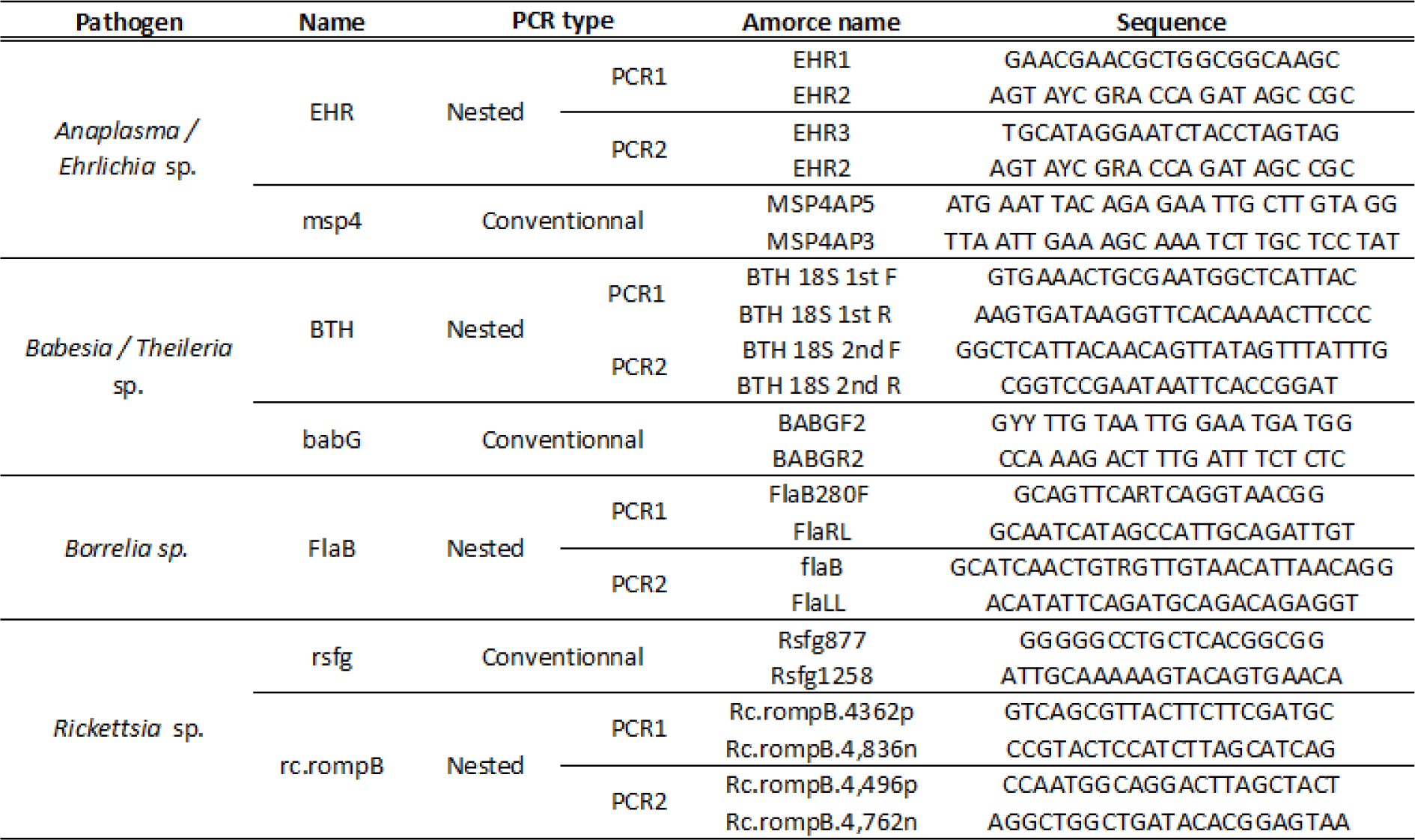
Primers used to confirm each pathogen detected by the microfluidic chip, through conventional and/or nested PCR.

Finally, a specific detection of CCHFV was performed on tick samples. All RNA extracts were screened for CCHFV by another RT-qPCR targeting a 3’ non-coding region of the CCHFV genome using specific primers and probes (Sas et al., 2018). rRT-PCR Taqman assays were performed in a final volume of 20 μL using the LightCycler 480 RNA Master Hydrolysis Probes Master Mix (Roche Applied Science, Germany) at 1 X final concentration, with 0.5 μM specific primers and 0.25 μM probes and 2 μL RNA. Negative (ultra-pure water) controls were included in each run. rRT-PCR thermal cycling conditions were as follows: 63°C for 3 min, 95°C for 30 s, 45 cycles at 95°C for 10 s then 60°C for 30 s, followed by cooling at 40°C for 10 s.

### Statistical analysis

The statistical analysis was conducted using R software (version 4.2.1, R Core Team).

In order to investigate the potential difference in the presence of *Rickettsia* between male and female ticks, a chi-square test was performed, as a relevant method commonly used to assess associations between categorical variables. The variables of interest were the presence of *Rickettsia* (1,0) and the gender (male or female), and the test was conducted to determine if there was a statistically significant association between gender and the presence of *Rickettsia*.

## Results

### 1. Identification of microorganisms detected in H. marginatum

DNA and RNA of ten of the targeted tick-borne microorganisms were identified in the 1,195 tick samples (973 individual ticks and 222 pools of 1 to 10 ticks). Using the microfluidic method on individual ticks, we detected 2 ticks positive for *Borrelia* spp, 3 for *A. marginale*, 33 for *A. phagocytophilum*, 201 for *Anaplasma* spp, 3 for *Ehrlichia* spp, 878 for *R. aeschlimannii*, 957 for *Rickettsia* spp, 8 for *B. henselae*, 4 for *Bartonella* spp., 68 for *Theileria* spp, 22 for *T. equi*, 13 for West Nile Virus, 1 for CLE and 1,160 for FLE. In terms of tick pools, the same method identified 19 positive pools for *Anaplasma* spp., 215 pools for *R. aeschlimannii*, 214 for *Rickettsia*. spp., 10 for *Theileria*. spp., 5 for *T. equi*, 216 for FLE and 3 for WNV. Except for the well-known CLE and FLE (and because the pathogenic forms *C. burnettii* and *F. tularensis* were never detected in this study), any other amplified microorganisms suspected to be pathogenic were sequenced from some positive samples to be confirmed.

For *Rickettsia* spp., most of samples amplified specifically for *R. aeschlimannii* were also amplified in *Rickettsia spp.*, using the microfluigdim method. 18 amplicons (5 combining the two targets, 3 amplifying only *R. aeschlimannii* and 10 amplifying only *Rickettsia* spp.) were sequenced. Of the 10 readable sequences, all sequences confirmed the presence of *R. aeschlimannii*. One of these sequences has been deposited in GenBank (AN: PP379722), showing 98.77% homology with a strain isolated from a *H. marginatum* found in England in 2019 (AN: MT365092) and 97.85% homology with another strain detected in Corsica in 2019 (AN: MK732478) in a *Rhipicephalus bursa* tick.

For *Anaplasma* spp., all samples amplifying *A. marginale* were also positive for *Anaplasma* spp., which was not necessarily the case for *A. phagocytophilum*, for which some samples were positive for *A. phagocytophilum* but not *Anaplasma* spp., and many samples were only amplified using primers targeting *Anaplasma* spp. Nineteen amplicons representative of the four cases (*A. marginale* + *Anaplasma* spp. / *A. phagocytophilum* only / *A. phagocytophilum* + *Anaplasma* spp. / *Anaplasma* spp. only / *A. phagocytophilum* only) were sequenced, and only confirmed samples that combined the amplification of both specific and generic targets. The samples positive only for *Anaplasma* spp. were finally identified as *Devosia* spp. by sequencing confirmation PCR products, which is an ubiquitous soil bacteria that is considered as a PCR cross contaminant. At the end, two *Anaplasma* species were confirmed: *A. phagocytophilum* and *A. marginale*. The sequence of *A. phagocytophilum* (AN: PP227262) showed 99% homology with the reference sequence from an *A. phagocytophilum* strain isolated in a *Rhipicephalus microplus* tick collected on a cow in Taiwan in 2021 (AN: OL690562.1) and 99.12% with a sequence from a strain isolated in an *Ixodes ricinus* tick, in Hérault, France in 2010 (AN : OR426543). For *A. marginale*, the sequence (AN : PP218689) showed 96% homology with a reference sequence from an *A. marginale* strain isolated in a *H. schulzei* tick collected in Iran in 2019 (AN: MK310488) and 95,27 % from a strain isolated in cattle in 2017 in Spain (AN: MT664983).

For *Ehrlichia* spp., two amplicons were sequenced and were confirmed as *E. minasensis*. The resulting sequence deposited in GenBank (AN: PP227261) showed 99% homology with a reference sequence (AN: NR*_148800.1)* of a strain that was isolated from the hemolymph of a *Rhipicephalus (Boophilus) microplus* tick from Brazil in 2019. Moreover, the sequence detected showed 98% homology with a strain (AN : JX629805) detected in a *Rhipicephalus microplus* tick from Brazil in 2016. The detected sequence in Corsica was 100% identical with the brazilian strain (Vincent Cicculli et al. 2019).

For *Theileria* spp., most of samples that were amplified specifically for *T. equi* were also amplified for *Theileria* spp., except two samples positive only for *T.equi*, and other samples were only amplified for *Theileria* spp. No sample was amplified for *T. annulata*. Sixty six amplicons combing the three cases (*T. equi* only / *T. equi + Theileria* spp. / *Theileria* spp. only) were sequenced and confirmed the presence of *T. equi* for samples amplified for both targets as well as those amplified only for *T. equi*. All amplicons for *Theileria* spp. only produced sequences that could not allow species-specific identification despite repeated sequencing (unreadable sequences). For *T. equi,* the same sequence (AN: PP227191) was obtained for all samples confirmed and showed 99% homology with a reference sequence from a *T. equi* strain isolated in Chile in 2020 (AN: MT463613.1) and 99,81% homology with a sequence from a *Rhipicephalus bursa* tick in Corsica in 2019 (AN: MK732476.1).

Nine ticks detected positive for *Theileria* spp. only were finally confirmed by sequencing as infected by *Babesia occultans*. The sequence deposited in GenBank (AN: PP227192) showed 99% homology with a sequence of *B. occultans* isolated from a *H. excavatum* tick collected on cattle in Turkey in 2022 (AN : OM066180.1).

One individual tick and one pool of ticks from Corsica were positive for the West Nile Virus. The amplification product was sequenced (AN: PP379723) and showed 100% homology with a reference sequence from the complete genome of West Nile Virus detected from a snowy owl in Germany in 2019 (AN : LR743458.1).

### 3. Proportions of microorganisms in H. marginatum, according to host origin and spatial distribution

Results of individual and pooling detections are presented in Table 3.

**Table 3:**
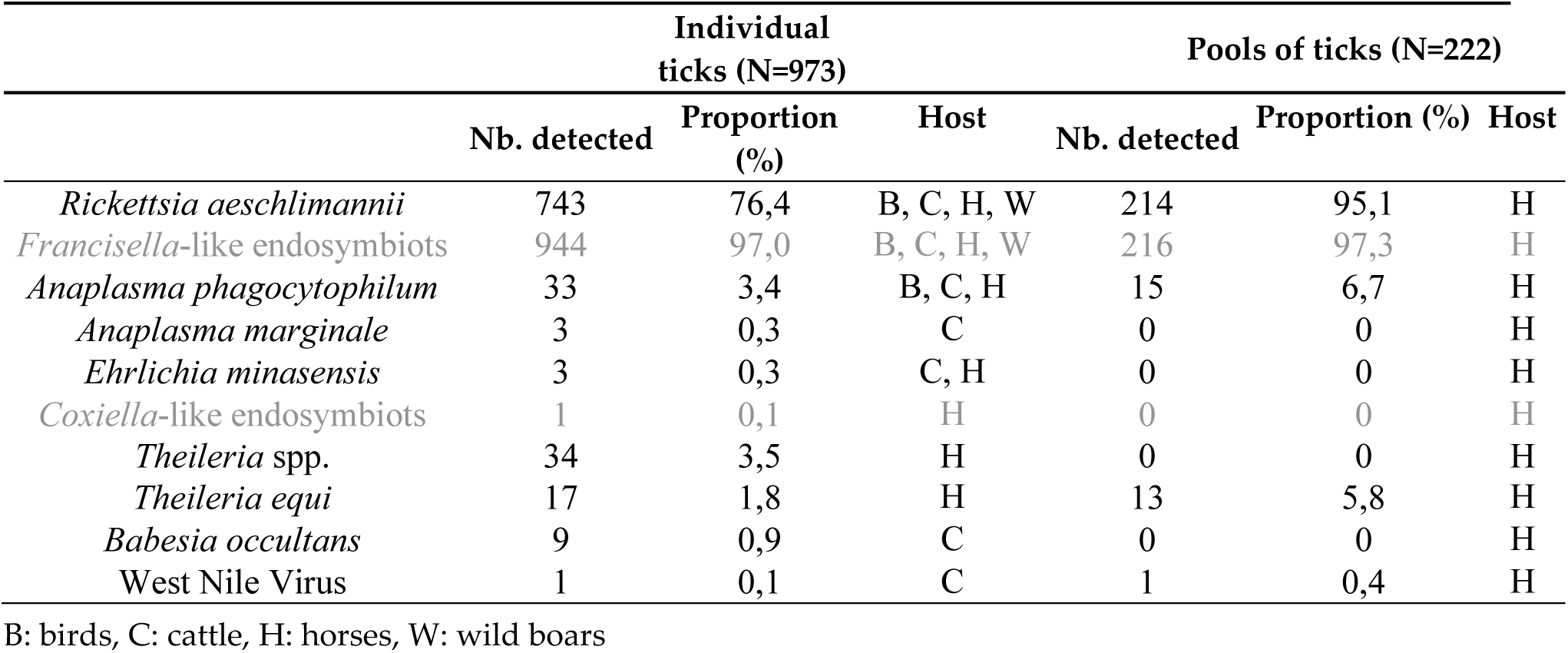
Microorganisms detected and confirmed in *H. marginatum* ticks, tested individually or by pools. The currently known or assumed pathogens are in black and the endosymbionts in grey.

*FLE* was the most prevalent microorganism, as detected in more than 95% of individual ticks and pools of ticks of *H. marginatum*. *R. aechlimannii* was detected in more than 75% of the analyzed individual ticks and 95% of pools tick of *H. marginatum*. Conversely, we only identified a CLE in one tick collected on a horse in Var. Then, proportions of infection due to all the other targeted tick-borne pathogens ranged from 0.1% to 3.5% (up to 6.7% in pools of ticks). If we remove the associations between *R. aeschlimannii* that was highly prevalent and the other pathogens, the likelihood of coinfections in *H. marginatum* ticks was very low. Only 3 coinfections were observed, two ticks carrying both *B. occultans* and *A. marginale* and one tick carrying both *E. minasensis* and *A. marginale*.

Although equine piroplasmosis is commonly reported in horses from the South of France (Rocafort-Ferrer et al. 2022) and our ticks were mainly collected on horses, *T. equi* was detected in a very small number of ticks, 17 individual ticks from mainland France and 13 pools of ticks from Corsica, all collected on horses. No *B. caballi* was detected in the analyzed ticks. *B. occultans* was detected in 9 individual ticks collected only on cattle in Corsica. For bacteria, *A. marginale* was only detected in ticks collected on cattle in Corsica, while *A. phagocytophilum* was mainly found in ticks collected from horses but also on cattle and to a lesser extent in ticks collected on birds (n = 2/37). Birds were mainly infected by *R. aeschlimannii* (n = 15/37). *E. minasensis* was detected in 3 ticks, two from Corsican cattle and one from a horse on the mainland. Only one virus, the West Nile virus, was detected in one tick collected from a cow and one pool of ticks collected from a horse, both in Corsica. Conversely, *H. marginatum* ticks were all negative for CCHFV. No specific pathogens, except *R. Aeschlimannii* (n = 6/7), were found in ticks collected on wild boar. Aside from minor differences between ticks collected on horses versus cattle, and considering the significant underrepresentation of hosts other than horses, which limited statistical comparisons, the panel of microorganisms appeared to be quite similar across the different hosts.

Finally, we did not observe significant differences in the presence of most of the detected microorganisms between male and female ticks (*A. phagocytophilum* : χ^2^ = 0.46 df = 1, *P* = 0.49; *A. marginale* : χ^2^ = 1.37, df = 1, *P* = 0.24; *E. minasensis* : χ^2^ = 1.37, df = 1, *P* = 0.24; *Theileria* spp. : χ^2^ = 1.37, df = 1, *P* = 0.24; *T. equi* : χ^2^= 0.001, df = 1, *P*= 0.97). However, *R. aeschlimannii* was significantly much more present in males than females (n(M) = 388; n(F) = 340; χ^2^ = 10.28, df = 1, *P* = 0.001) and *B. occultans* in females than males (n(M) = 1; n(F) = 8; χ^2^ = 4.14, df = 1, *P* = 0.04).

## Discussion

In the present study, several tick-borne microorganisms, including both pathogens and endosymbionts, were identified in *H. marginatum* ticks primarily collected on horses, with some samples taken from other animal species, on the island of Corsica as well as on mainland France, where this tick species has recently settled (Vial et al., 2016). The microfluidic method allowed the detection of eight known or suspected genera and species of pathogens, specifically *R. aeschlimannii*, *A. phagocytophilum*, *A. marginale*, *E. minasensis*, *Theileria* spp., *T. equi*, *B. occultans* and West Nile Virus, along with two endosymbionts species (FLE and CLE), all of which being already demonstrated to circulate in France. The most prevalent microorganisms detected were FLE and *R. aechlimannii* with a large proportion of ticks infected, while the infection rates for the other microorganisms were relatively low, ranging from 0.1 to 6.7%, depending on whether individual or pooled ticks were analysed.

The microfluidic method demonstrated significant potential for high-throughput, targeted detection of multiple parasites, bacteria and viruses, simultaneously, using only a minimal amount of material enabling detection at the individual level. However, this approach may overlook unexpected or new microorganisms present in the sample but that were not included in the pre-defined list of targeted candidates. This limitation raises concerns when investigating the panel of microorganisms that could be introduced by an invasive exotic tick species such as *H. marginatum*. Consequently, it was necessary to develop and adapt the microfluidic array specifically for our study to comprehensively identify the diversity of microorganisms that could be transmitted by, or at least detected in, such invasive exotic tick species within both its native and expension geographical zones. Furthermore, our strategy to target not only specific microorganism species but also broader genera, and to confirm the targets by sequencing, enhanced our detection and identification capacities. For instance, *B. occultans* was subsequently confirmed in samples that were initially positive for *Theileria* spp. However, for efficiency reasons, confirmations using sequencing were performed only on a limited subset of detected samples, resulting in unavoidable extrapolations for samples exhibiting similar detection patterns. This is illustrated by *Anaplasma* spp. and *A. phagocytophilum*, for which out of 19 samples sent for sequencing, two sequences confirmed the presence of *A. phagocytophilum*, while the remaining sequences corresponded to *Devosia* spp., an ubiquitous soil bacterium contaminating numerous samples and co-amplified by our PCR primers designed for *Anaplasma* spp. From this observation, the samples that tested positive for *A. phagocytophilum* on the microfluidic chip and exhibited the same profile as the two samples with confirmed sequences, were all classified as *A. phagocytophylum*. Conversely, the samples amplifying *Anaplasma* spp. only were ultimately deemed negative for *Anaplasma*, as we assumed the false positive results were due to contamination by *Devosia*. This may be a limitation of the microfluigdim method, which avoids to investigate intraspecific diversity within detected pathogens, but this was clearly not the primary aim of our study. Instead, this high-throughput approach allowed us, for the first time, to provide a comprehensive panel of microorganisms encountered in 973 individual *H. marginatum* ticks and 900 ticks grouped in 222 spools collected across 9 French departments, from Spanish to Italian borders and including Corsica island. Despite potential biases related to differing sampling efforts in Corsica island and mainland France, this method also enabled the first prevalence estimates of microorganisms (including TBPs) carried by *H. marginatum,* based on their geographic location and animal hosts. By integrating ecological and epidemiological data on tick-host-pathogen associations, our study generated first hypotheses regarding the potential role of *H. marginatum* as vector for the detected pathogens. However, since *H. marginatum* is a hunting tick, all samples in our study were collected from animals during their blood feeding. This raises questions about the origin of microorganisms’ DNA or RNA detected in ticks; specifically, whether the ticks were truly infected or merely carried microrganisms from their infected animal hosts via the remaining blood meal in the tick midgut. While experimental infection studies are scarce, they are crucial for confirmainng a tick species’ vector competence, namely its ability of to acquire a pathogen from an infected host, replicate and maintain it, and transmit it to a new, naïve host.

Initially, the tick sampling was included in a monitoring program for emergence risk of CCHF, under the auspices of Ministère de l’Agriculture, de la Souveraineté alimentaire et de la Forêt. CCHFV was not detected in any of *H. marginatum* tested in our study, despite serological evidence of antibodies against the virus in cattle from the same areas where the sampled horses were located, which were infested with *H. marginatum* (Grech-Angelini et al. 2020). Confirmation of these serological results using the pseudoplaque neutralization test (PPNT) on CCHFV cultures in BSL-4 conditions suggests that the virus, or an antigenically similar variant (CCHFV-like), circulates in the natural enzootic cycle between ticks and non-human vertebrate hosts (Grech-Angelini et al. 2020). In Europe, CCHFV infection prevalence in ticks ranges from 0.5% to 3.70% (Cicculli et al., 2022), so our sampling effort may have been insufficient to detect at least one infected tick. Most of the ticks collected between 2016 and 2020 were collected from horses, and based on studies of *H. marginatum* hosts and their abundance in France, as well as their varying ability to replicate CCHFV and infect new naïve ticks during blood feeding, most French *H. marginatum* are likely to be in contact with "poor" hosts, such as birds or horses, which have a low or negligible probability of transmitting the virus (Bernard et al. 2022). Another hypothesis could be the existence of highly localized CCHFV transmission hotspots, which may not have been sampled due to difficult access during the optimal virus’ circulation period, an issue that requires further investigation. This hypothesis was supported by the first detection of CCHFV in *H. marginatum* ticks in France in 2023, where targeted sampling was conducted (Bernard et al., 2024), unlike the more general sampling in this study. Additionally, the lack of detection of CCHFV in our study could be attributed to a too low virus load in the ticks to be detected or to a fast degradation of the genetic material (RNA virus), despite freezing of ticks at −80°C after morphological identification (Cicculli et al. 2022).

Actually, the only virus detected in the tick samples was WNV, in one individual tick, which was collected from a Corsican cow, and in one pool of ticks from a Corsican horse. Since cattle are not involved in the natural cycle of WNV, it is likely that the tick became infected during a previous blood meal, at immature stage on a bird. Birds play a key role in the WNV cycle and are considered as reservoirs of the virus, as they are able to reinfect vectors, typically mosquitoes, and possibly ticks that feed on them (Vaughan, Newman, et Turell 2022). Regarding the tick pool of 10 males collected from a horse, it is important to note that horses are epidemiological dead-ends for WNV, as they do not produce a sufficient viremia to infect vectors, including ticks despite larger volumes of blood ingested (David et Abraham 2016). Additionally, male ticks engorge much less than females even not at all, making it unlikely that the virus was transmitted from the horse to the ticks. This raises the question of *H. marginatum* being able at least of transstadial transmission of WNV, although this alone does not confirm full vector competence. However, horizontal transmission of WNV by *H. marginatum* to an uninfected vertebrate host, in both immature and adult stages, has been experimentally demonstrated in Portugal (Formosinho et Santos-Silva 2006). Given that WNV can cause neurological disease and death in humans, our findings highlight the need to intensify monitoring of WNV in *H. marginatum* and further innvestigate the potential of this tick species to transmit the virus.

Concerning *Francisella* and *Coxiella* detected in our study, they were considered as *Francisella*-like and *Coxiella*-like non-pathogenic mutualistic and/or commensal endosymbionts, as previously described (Buysse et al. 2021; Hussain et al. 2022). In our study, about 97% of *H. marginatum* were infected by FLE, and only one tick was positive for CLE, which is coherent with previous observations that FLE outcompetes CLE in ixodid ticks (Kumar et al. 2022). High prevalence of FLE in *H. marginatum* ticks was previously observed, as it seems to be an ancestral primary endosymbiotic organism (mandatory to tick survival), being involved in nutritional symbiotic interactions including B vitamins biosynthesis pathways critical for tick metabolism (vitamin B9 ( folate), B2 (riboflavin) and B8 (biotin)) (Buysse et al. 2021). B vitamins are present at low concentrations in the blood of ticks’ hosts compared to the amount required for arthropods development (Zhong, Jasinskas, et Barbour 2007; Duron et al. 2018; Serrato-Salas et Gendrin 2023), resulting in the need of production by bacterial endosymbionts. Deprivation of their nutritional symbionts induced in ticks a complete stop of growth and moulting together with lethal physical abnormalities, which can be fully restored by exogenous supplementation with B vitamins(Buysse et al. 2021; Cibichakravarthy et al. 2024).

The second most abundant bacterium identified in *H. marginatum* ticks was *R. aeschlimannii,* detected in 75% of individual ticks and 95% of tick pools, and thus in much larger proportions than any other bacteria detected in our study. *R. aeschlimannii* was first isolated from *H. marginatum* ticks in Morocco in 1997 (Beati et al. 1997), and is now regularly reported in ticks across the Mediterranean region (Parola et al. 2014). In Corsica, Grech-Angelini et al (2020) detected *R. aeschlimannii* in 100% of the *H. marginatum* tick pools they tested. Pooling ticks can inflate infection prevalence estimates, as a pool tests positive even if only one tick within it is infected. This was evident in our study, where pooled tick samples showed more frequent positive results compared to individual ticks, with only two negative pools, both containing a single tick. Given the high prevalence of *R. aeschlimannii* in ticks from both mainland France and Corsica, and considering its potential pathogenicity in humans, the risk of human exposure to this bacterium likely exists, even though humans are not common hosts for *H. marginatum*. In Corsica, it has been hypothesized that the high presence of this bacterium may be correlated to human cases of tick-borne spotted fever reported on the island (Grech-Angelini et al. 2020). It was previously reported humans cases of spotted fever in North Africa and South Africa caused by *R. aeschlimannii* infections, and in 2010 a first case occurred in Southern Europe, in a Greek patient (Germanakis et al. 2013), followed by two addditional cases in Italy in 2016 and in Germany in 2019 (Tosoni et al. 2016; « Cas suspect de fièvre tachetée à Rickettsia aeschlimannii en Allemagne », s. d.). Notably, the strain detected in these cases shares 100% homology with the strain identified in the Greek patient (GeneBank Accession Number: JF803904.1). However, no human cases of tick-borne spotted fever due to *R. aeschlimannii* have been reported so far on the French mainland, raising questions about the pathogenic role of *R. aeschlimanii* found in *H. marginatum* ticks. *R. aeschlimannii* can be transmitted transovarially from *H. marginatum* females to their offspring, supporting the idea that *H. marginatum* could act as a reservoir for this bacterium (Matsumoto et al. 2004; Azagi et al. 2017). Given the significantly higher prevalence of *R. aeschlimannii* compared to other pathogenic bacteria in *H. marginatum*, an alternative hypothesis suggests that it may function as a secondary symbiont within these ticks (Joly-Kukla et al., 2024). Similar to *Rickettsia peacockii* in *Dermacentor andersoni*, secondary symbionts can regulate the development of other infectious *Rickettsia* (Baldridge et al. 2004). While primary symbionts are virtually fixed in the tick population because vital, secondary symbionts, though non-essential, may provide ticks with a selective advantage. At this point, it is unclear whether *R. aeschlimannii* acts as a pathogen, a symbiont, or both. Further investigations are needed, including its specific detection in tick salivary glands to confirm its ability for horizontal transmition through tick bite. Additionally, experimental trials involving the removal of *Rickttsia* from ticks would help clarify its role in tick metabolism and development.

Apart from *R. aeschlimannii*, bacterial DNA detected in *H. marginatum* was mostly represented by the genus *Anaplasma*, namely *A. phagocytophilum* and *A. marginale, remaining* both at low levels (3,7% in individual ticks, 6,7 % in tick pools for *A. phagocytophilum* and less than 1% in individual ticks (not detected in tick pools) for *A. marginale*). *A. phagocytophilum* is a zoonotic obligate intracellular bacterium causing Tick-Borne Fever in cattle and sheep, Human Granulocytic Anaplasmosis (HGA) in humans, and Equine Ehrlichiosis (Rikihisa 2011), and *A. marginale* is responsible for the erythrocytic bovine anaplasmosis that can affect various species of domestic and wild ruminants, including cattle, and that is largely distributed in France (Aubry et Geale 2011). A recent study showed that in the French Pays Basque, 22.4% of collected *Rhipicephalus bursa* ticks contained *A. phagocytophilum* DNA (Dahmani et al. 2017), indicating other tick species more adapted to carry *A. phagocytophilum*. This TBP has been reported in several European countries (Matsumoto et al. 2006). HGA cases have been documented in mainland France, with a first case identified in 2003 and others reported sporadically since then (Edouard et al. 2012). Human cases have also been recorded in continental Italy (Ruscio et Cinco 2003). This is not the first time *A. phagocytophilum* has been detected in *H. marginatum*, as one infected tick was found in Israel in 2007 (Keysary et al. 2007), and three in Tunisia in 2012 (M’ghirbi et al. 2012). As observed in our study, the prevalence of *A. phagocytophilum* infection in ticks has consistently been low, despite its high presence in cattle (Woldehiwet 2006) and horses, with 67% of horses testing positive (M’ghirbi et al. 2012). These animals likely serve as hosts for the adult stages of *H. marginatum*. In France, *A. phagocytophilum* is present in about 22% of rodent species across different regions (Chastagner et al., 2016), 0.6% of cattle, 0.4% of birds and 3.3% of common blackbirds (Rouxel et al., 2024), but 0% of horses studied (Dugat et al., 2017). Under these conditions, the prevalence of infection in ticks carrying feeding on infected hosts would not increase significantly. Additionally, our detection of the bacterium in non-engorged adult ticks raises questions about their potential role as reservoirs and vectors for *A. phagocytophilum*, especially given that the hosts of the tick’s immature and adult stages (birds, cattle, horses) are susceptible to this bacterium and that transovarial transmission occurs in other tick species, what desserves further studies on the vector competence of *H. marginatum* for *Anaplasma* bacteria. Additionally to *A. phagocytophilum*, *A. marginale* DNA was found in 3 *H. marginatum* ticks collected from cattle in Corsica, which is consistent with previous reports in ticks from Corsica (Dahmani et al. 2017; V. Cicculli et al. 2019), including *H. marginatum* (Grech-Angelini et al. 2020). The primary vectors of this bacterium are *Rhipicephalus* spp. and *Dermacentor* spp. found in tropical and subtropical regions but also in Europe. The activity periods of these ticks may overlap with those of *H. marginatu*m, suggesting the tick could become infected while feeding on an infected animal. The bacterium tipically circulates in late spring and early summer, which coincide with *H. marginatum*’s period of activity, as well as in autumn. Although not considered as a primary vector of this bacterium (Atif 2015), our findings indicate that the presence of this pathogen in our ticks likely reflects their host’s infection, as we only detected infected ticks from cattle.

Another bacterium identified in *H. marginatum* was *E. minasensis*, a recent member of the genus *Ehrlichia* and responsible for bovine Ehrliciosis. In 2014, it was shown to naturally infect cattle in Brazil and cause clinical Ehrlichiosis in an experimentally infected calf (Aguiar et al. 2014). The distribution of *E. minasensis* extends beyond the Americas (Gajadhar et al. 2010; Cruz et al. 2012; Aguiar et al. 2014), with reports of the bacterium in Pakistan (Rehman et al. 2019), Ethiopia (Hailemariam et al. 2017), South Africa (Iweriebor et al. 2017), Israel (Thomson et al. 2018), and it was recently detected in Corsica island in 3.7% of *H. marginatum* collected from cattle (Vincent Cicculli et al. 2019). Although the vector competence of ticks for this bacterium has yet to be assessed, *E. minasensis* has already been identified in several tick species including *R. microplus* (Cruz et al. 2012; Carvalho et al. 2016), *R. appendiculatus* (Iweriebor et al. 2017) and *H. anatolicum* (Rehman et al. 2019). In addition to infecting bovines (Gajadhar et al. 2010; Aguiar et al. 2014; Hailemariam et al. 2017), *E. minasensis* infects also cervids (Lobanov et al. 2012), dogs (Thomson et al. 2018), and horses (Muraro et al. 2021). In our study, *E. minasensis* was detected in three ticks, two from cattle in Corsica and one from a horse in mainland France. All three ticks were fully engorged and still attached to their host when collected, suggesting this detection may simply be due to their feeding on infected animals. However, we cannot rule out the potetial vector competence of *H. marginatum* for this newly described pathogen, which requires further investigation.

Concerning protozoans, DNA detected in *H. marginatum* was mostly represented by the genus *Theileria*. This was *T. equi*, one of the causative agents of Equine Piroplasmosis (EP) in France, which can also be due to *B. caballi* (Wise et al. 2013) that was not detected in *H. marginatum* in our study raising questions on whether this pathogen is not or no longer circulating in horses in our study area in South of France, which has been previously observed (Rocafort-Ferrer et al. 2022), or that the primers used for PCR amplification also failed to detect *B. caballi*, which desserves futher experiments on the methods to detect the diversity of piroplasms circulating in France. *T. equi* infects wild and domestic equines worldwide, and are enzootic in many countries including several European countries bordering the Mediterranean sea such as France (Tirosh-Levy et al. 2020). This parasite is transmitted by hard tick species belonging to several genera, including *Dermacentor*, *Rhipicephalus*, *Amblyomma*, *Hyalomma*, and *Haemaphysalis* (Scoles and Ueti 2015; Tirosh-Levy et al. 2020). This is not the first instance of this parasite beeing identified in *H. marginatum* in France; a previous study reported that 43% of *H. marginatum* ticks collected from horses in Camargue tested positive for the parasite in 2022 (Rocafort-Ferrer et al. 2022). Notably, our infection rate was considerably lower than that observed in this study. This significant discrepancy may be attributed to the specific design of the tick collection, which focused on stables where cases of piroplasmosis were frequently reported for Rocafort-Ferrer et al. versus stables of all kinds in our study. EP caused by *T. equi* is known to be widespread in France, particularly in the southern regions, with many asymptomatic carrier horses (Rocafort-Ferrer et al. 2022). Although the horses in this study were asymptomatic, a high proportion of horses in the same study area had antibodies against *T. equi* (Guidi et al. 2014), indicating that the parasite circulates in the area. But this is opposite to the low proportion of ticks positive for *T. equi* we found. This is all the more surprising since, once infected with *T. equi*, horses can remain infected for the rest of their lives, becoming asymptomatic carriers of the parasite, and thus constituting a source of infection for uninfected ticks (Friedhoff et al. 1990). Our approach has highlighted the likely underestimation of the infection rate, as we used primers that did not allow the detection of all genotypes potentially circulating in France. In fact, Joly-Kukla et al. (2024) used the same method and later re-evaluated *T. equi* infection rates by targetting a different gene (18S rRNA) through qPCR, which can detect multiple *T. equi* genotypes. Furthermore, our findings may be influenced by the possibility that the areas sampled in our study do not have high levels of parasite circulation, contrary to Camargue (Rocafort-Ferrer et al. 2022). Certain tick species, such as *Rhipicephalus turanicus*, which are present in Camargue but less common in other parts of southern France, are known vectors of the parasites responsible for EP. This could account for a higher local circulation of the parasite and, consequently, a greater prevalence of infection in ticks. However, this is not an isolated instance; previous studies have also shown cases where the prevalence of infection in horses was high although the infection rate in ticks remained low. For example, Nadal (2019) reported that where 50% of horses were infected, yet only 3% of the ticks tested positive, questionning whether an insufficient parasite load to be infectious through blood meal or a moderate role of reservoir played by ticks. This could also be linked to our observation that about 5% of our collected ticks tested positive for *Theileria* only at genus level, suggesting a low circulation in horses and therefore a very low transmission to ticks, avoiding precise identification of the parasite species because of insufficient molecular material. While concerns were raised about the risk of introducing pathogens belonging to the genus *Theileria* causing Tropical Theileriosis with the arrival of *H. marginatum* in France, our study found no*T. annulata* DNA in the collected ticks, neither did we detect *T. buffeli*, a benign cattle pathogen previously detected in Corsica (ICTTD, 2000). Interstingly, in about two-thirds of the cases where only *Theileria* spp. was identified, we were not able to determine the specific species of *Theileria* because of unsuccessful sequencing. Several hypotheses can be considered for these cases. For ticks collected from cattle, we can wonder if it is *T. buffeli* that is detected, given its presence in cattle, even though no species other than *T. equi* was characterized. Alternatively, it may be *T. equi*, as our primers may not have been suitable for detecting all genotypes circulating in France, which requires the definition of new primers to improve the detection of all *Theileria* species potentially transmitted by ticks..

We have finally detected *B. occultans* by sequencing in nine cattle ticks from Corsica amplifying first *Theileria* spp. with the micro array. Some studies conducted in Turkey and Tunisia, have previously detected *B. occultans* in *H. rufipes*, *H. marginatum*, *R. turanicus*, and *H. excavatum* ticks (Aktas, Vatansever, et Ozubek 2014; Ros-García et al. 2011; Orkun 2019; Orkun et al. 2020). So far, only *H. rufipes* has been confirmed as a vector of *B. occultans* (Ozubek et al. 2020) and *H. marginatum* is not described as a potential vector of this pathogen. In our study, we can assume that ticks engorged on cows carrying this parasite and got infected during their blood feeding. Nevertheless, it cannot be ruled out that an evolutive adaptation could lead to *H. marginatum*-borne transmission of *B. occultans* in southern Europe. This deserves further investigations in all tick species collected from cattle populations, as recently done in Morocco (Elhachimi et al. 2021).

Except for known endosymbionts, and the suspected endosymbiotic status of *R. aeschlimanii*, our study revealed that the infection prevalence of endemic pathogens remains very low in the tick *H. marginatum*, settled in the south of France since 2016. The vector competence of this tick species for *T.equi* and *A. phagocytophilum* is still uncertain, but it has to be considered as it is almost demonstrated for CCHFV (Kiwan 2024, personal communication), confirming the risk of emergence of *Hyalomma marginatum*-borne pathogens. Additionally, we did not identify any new exotic pathogens that could have have been introduced by *H. marginatum* from its original distribution area. Based on this information, and considering that the epidemiological situation is dynamic and may evolve, we propose that *H. marginatum* serves as valuable sentinel arthropod for anticipating significant health risks for humans or animals in France. Therefore, it warrants ongoing monitoring due to its potential to spread on long distances through bird migrations and terrestrial animal trade.

## Acknowledgements

We would like to thank all the people who were involved in the field tick collections. We would like to thank Maxime Duhayon and Karine Huber for technical support in managing tick processing and biological material extraction. We would like to thank all cattle and horse owners for collaborating with us on this study.

## Funding

The authors thank the funders who made this work possible: French Ministry of Agriculture—General Directorate for Food (DGAl, grant agreement: SPA17 number 0079-E), European Funds for Regional Development (FEDER, Grand-Est), French Establishment for Fighting Zoonoses (ELIZ) and the Association Nationale Recherche Technologie (ANRT, grant agreement number: 2019-1145), Défi clé RIVOC (Occitanie Region): Holis-tiques project. This research was funded by grant from the French Agency for Food, Environmental and Occupational Health & Safety (ANSES).

## Conflict of interest disclosure

The authors of this preprint declare that they have no financial conflict of interest with the content of this article. Thomas Pollet, Sara Moutailler and Laurence Vial are recommenders for PCI infections.

## Author contributions

Conceptualization: CB, MB, BC, LV, PH; Data curation: CB, CG, CJK, VC, SGA, PEP, SM, PH, LV; Funding acquisition: MB, BC, SM, PH, LV;; Methodology: CB, CG, SGA, SM, PH, LV; Project administration: CB, TP, MB, BC, SM, PH, LV; Supervision: CB, TP, MB, BC, SM, PH, LV; Validation: all authors; Writing – original draft: CB, PH, LV; Writing – review and editing : all authors.

## Data, scripts and codes availability

Data are available online: https://gitlab.cirad.fr/astre/data_tickpathogens_cbernard2025

## Notes

https://gitlab.cirad.fr/astre/data_tickpathogens_cbernard2025

### Competing Interest Statement

The authors have declared no competing interest.

### Summary of Updates

Manuscript updated to be submitted to PCI infections

